# Computational microbiology of soil organic matter mineralization: Use of the concept of curve skeleton to partition the 3D pore space in computed tomography images

**DOI:** 10.1101/2024.10.24.620029

**Authors:** Zakaria Belghali, Olivier Monga, Mouad Klai, El Hassan Abdelwahed, Lucie Druoton, Valérie Pot, Philippe C. Baveye

## Abstract

Recent advances in 3D X-ray Computed Tomography (CT) sensors have stimulated research efforts to unveil the extremely complex micro-scale processes that control the activity of soil microorganisms. Classical methods for the numerical simulation of biological dynamics using meshes of voxels, such as the Lattice Boltzmann Method (LBM), tend to require long computation times. The use of more compact geometrical representations of the pore space can drastically decrease the computational cost of simulations. Recent research has introduced basic analytic volume primitives to define piece-wise approximations of the pore space to simulate drainage, diffusion, and microbial mineralization of organic matter in soils. Such approaches work well but a drawback is that they give rise to non-negligible approximation errors. In the present article, another alternative is proposed, where pore space is described by means of geometrically relevant connected subsets of voxels (regions) regrouped on the basis of the curve linear skeleton (3D medial axis). This curve skeleton has been adopted to characterize 3D shapes in various fields (e.g., medical imaging, material sciences, etc.) but the few publications that have used it in the context of soils, have dealt exclusively with the determination of pore throats. This technique is used mostly to describe shape and not to partition it into connected subsets. Here, the pore space is partitioned by using the branches of the curve skeleton, then an attributed relational graph is created in order to simulate numerically the microbial mineralization of organic matter, including the diffusion of by-products. This new representation can be used for graph-based simulations, which are different from voxel-based simulations.

## 1 Introduction

Soils worldwide contain a very large stock of organic matter, whose mineralization by microorganisms has the potential to have a sizeable impact, positive or negative as the case may be, on the amount of greenhouse gases released to the atmosphere, and therefore on climate change [1-4]. Over the last four decades, the importance of soils in that context has prompted a very significant research effort on the description and prediction of the fate of organic matter in subsurface environments. Initially, the modelling component of that effort focused on the development of black-box models like Century or RothC, in which the metabolic activity of microorganisms was described simply via first-order kinetic equations, without a detailed accounting of the growth and spatial distribution of the biomass [5,6]. After the turn of the century, it became clear that much more attention needed to be paid to where in the soil pores microorganisms were located relative to the organic matter they fed on [7,8]. Fortunately, concomitant technical advances in X-ray computed tomography (CT) enabled researchers to routinely obtain 3-dimensional greyscale pictures of soils at progressively increasing resolutions, from which, after the development of suitable thresholding algorithms [9,10], binary images could be obtained, providing microscale information on the geometry and topology of soil pores in which microorganisms reside. Over the last decade, this information, complemented by a range of other microscale microscopic and spectroscopic measurements [11-13], has stimulated extensive research on the microscale modeling of soil microbial processes [14-17].

One of key hurdles that researchers have faced in the computational microbiology of soil organic matter mineralization is related to the sheer size of the image files that result from CT scans, with numbers of voxels in the hundreds of millions and upward for the highest attainable resolutions. The Lattice Boltzmann Method (LBM), which is generally the method of choice to describe the movement of water and the diffusion of nutrients in the pore space of soils, is very accurate [18] but requires computers with very large memories and is very demanding in CPU time with currently available computer technologies. In order to decrease the computational cost, the use of more compact geometrical representations of the pore space appears to be a valuable alternative. In the last decade, efforts have been devoted to the development of advanced computer vision and computational geometry tools in order to obtain better, less memory-intensive representations of the pore space [19-21]. These tools aim to provide compact, intrinsic, and relevant geometrical representations from the original set of voxels, which can lead to a considerable speeding up of the numerical simulation algorithms. Unlike older pore network models, which are based on an idealized pore space representation, the more recent methods take into account the exact geometry of the pore space as revealed by 3D CT images. Nevertheless, a drawback of approximation methods using geometric primitives is that, to be computationally advantageous relative to the LBM, they provide only a somewhat crude representation of the convoluted geometry of the pore space, which cannot be fully represented due to the comparative simplicity and compactness of spheres, for example. One of the main problems is the loss of voxels at the edges of the pore space.

In this general context, the present article introduces a novel approach in order, at the same time, to simplify the computational representation of the geometry of the pore space in CT images of soils and to make it less approximate. This approach is based, not on geometrical primitives, but on the partitioning of the soil pore space into connected subsets of voxels computed on the basis of the curve skeleton. This representation of the pore space, which ensures the inclusion of all voxels present in the pore space [22], is then used to model the microbial mineralization of organic matter in soils, paralleling earlier work [23,24]. Subsequently, following the methodology described in [25], we compare the modelling results obtained via this new approach with those produced by the classical Lattice-Boltzmann method. Lastly, we discuss future avenues for research.

## 2 Background and literature review

### 2.1) Skeletonization

In two dimensions, the so-called medial axis of a shape is defined as the locus of the centers of maximal inscribed disks. By definition, a disk is maximal if it is not strictly included in another disk inside the shape. In three dimensions, things are a little more complicated. The medial surface corresponds to the center of maximal inscribed balls, which in addition to a set of straight line segments or curves, also contains surfaces. This medial surface is often referred to as the surface skeleton and the process of obtaining it is termed the “skeletonization” of a geometrical shape [26].

These different concepts have been widely used in computational geometry since Blum’s landmark paper [27], and one can find in the literature various ways to compute the surface skeleton, depending on the initial description of the shape: triangular mesh, subset of voxels in a 3D image, point cloud sampling the boundary of the shape and so on. For instance, regarding shapes represented with a set of voxels in a 3D image, Xia and Tucker [28] computed a distance map to the shape boundary by solving the Eikonal equation with the Fast-Marching method. This distance map gives, for each voxel in the shape, the geodesic distance (i.e., the length of the shortest path inside the shape) to the boundary of the shape. The voxels of the surface skeleton are then extracted with the Laplacian of this map. Melkemi [29] and Amenta et al. [30] describe the Power Crust algorithm, which approaches the medial axis of a point cloud sampling the shape boundary. The surface skeleton can also be approximated using the 3D Delaunay Triangulation of a finite set of points forming a discretization (sampling) of the shape boundary. The set of the centers of the “Delaunay” spheres circumscribed to the Delaunay tetrahedra, and included within the shape, gives an approximation of the skeleton [22,31].

The sensitivity of the surface skeleton to little changes in the shape boundary is a drawback for several applications. There has been a tendency in the literature to invoke more robust, less sensitive approaches to the surface skeleton to address various tasks such as animation, motion tracking, shape recognition and analysis. When the details of the selected shape are irrelevant, the surface skeleton is simplified using the hierarchical removal of small maximal spheres. Such strategies are related to the notion of λ-skeleton [32], which presents undeniable interest in a number of contexts. However, in the type of situation that concern us, small maximal spheres can be filled with water, and thus diffusion processes can take place in such pores. Therefore, one should exercise caution when simplifying the computation of the surface skeleton in porous media and soils in particular.

Although the surface skeleton provides a sound foundation for skeletonization efforts, experience has shown that another type of skeleton, the curve skeleton, which is a 1D manifold rather than a 2D one, is much more straightforward to use for the representation of geometrical shape [33-35]. Several definitions have been proposed for this second type of skeleton, which is alternatively referred to as a “curve” or “curvilinear” skeleton in the literature. It can be defined as a 1-dimensional subset of the surface skeleton, fulfilling a supplementary condition, like the singularity of the Hessian of the distance map. There are other definitions based on morphological operators, like erosion and homotopic thinning.

Several discrete or continuous curve skeleton extraction methods have been proposed in the fields of discrete geometry and shape analysis. To this end, [36] describes a “homotopic thinning” algorithm. Zwettler et al. [37] adapted this algorithm in the specific case of tubular surfaces representing blood vessels for medical diagnostics. Sobiecki et al. [38,39] compared different methods for computing curve skeletons. They give a comparison of voxel-based methods with mesh contraction-based methods. Both are controlled by geometrical criteria such as homotopy, thickness, and preservation of details.

The use of the curve skeleton to segment 3D shapes is easy to illustrate and implement, but it has been explored only recently. Reniers and Telea [40] segmented the shape according to the critical points of the curve skeleton. The critical points were defined as triple cross road points, i.e., belonging to at least three branches of the curve skeleton. The curve skeleton is then segmented using the geodesic distances (length of the shortest paths inside the volume) of the critical points to the boundaries. Following the same principle, Brunner and Brunnett [41] presented a mesh partitioning algorithm combining voxelization and homotopic thinning. On that basis, they considered shapes for which the curve skeleton is relatively simple. However, that is generally not the case in soils.

### 2.2) Pore space geometrical modelling

In the earth sciences, Lindquist and Venkatarangan [42] and Lindquist et al. [43] use the medial axis in order to analyze the spatial distribution of pores in geological materials. Using this approach, Youssef et al. [44] address the quantitative 3D characterization of the pore space of real rocks within the context of petroleum extraction. The authors compare the results of the measurements with those of simulations for various porous media like sands and carbonates.

### 2.3 ) Analytic representation

Another alternative consists of defining an intrinsic analytic piece-wise approximation of the pore space using basic volume primitives. A drawback of most analytic approaches lays in the a priori choice of given geometric primitives. Within a pore network modelling context, a well-known algorithm is the Maximal Ball algorithm [45]. It consists of identifying the maximal spheres contained in the pore space and then approximating the volume by a subset of the spheres. This algorithm has been modified by Dong and Blunt [46], who propose a faster way to determine the set of maximal balls. In their approach, the network representing the pore space is composed of balls and cylinders associated with pores and throats. The biological simulation is then performed using this network. An alternative is to approximate the pore space with an optimal subset of maximal spheres and cylinders [19,31]. This optimal set is derived from the minimum set of balls (in a cardinal sense) recovering the λ-skeleton. Afterward, a numerical simulation of microbial mineralization is performed using a ball network [47]. Recently, several authors have proposed using more sophisticated geometrical primitives such as ellipsoids and generalized cylinders. For instance, the algorithm described in Kemgue and Monga [48] includes two steps. First, maximal spheres are clustered using the k-means partitioning method. Afterward, each cluster is approximated with a primitive. Finally, an optimal region-growing stage allows one to reduce the number of primitives. Regarding air-water interface extraction, a pore space representation with spheres or ellipsoids gives good results. Within other application contexts, including in medicine, several publications also deal with complex volume shape modelling using sophisticated primitives like for ellipsoids [22, 49-52]. Unfortunately, experience shows that these approaches cannot easily be extended to the pore space geometrical modelling in soils due to the higher shape complexity in these systems.

## 3 Results and Discussion

### 3.1) Geometrical modelling of the pore space

The geometrical modelling of the pore space using the novel approach based on the curve skeleton was very fast for each of the 4 images considered, when performed on a regular midrange laptop computer (AMD Ryzen 5 5500U, 8 Gigabytes RAM). The different computing times for the 4 data sets were, respectively, 61.23 seconds (data set 1, sandy loam soil), 124.47 seconds (data set 2, Fontainebleau quartz high porosity), 101.68 seconds (data set 3; Fontainebleau quartz low porosity), and 48.33 seconds (dataset 4, porous polycrystalline diamond). In terms of the associated demand of CPU time, one should note that this geometrical modelling has to be performed only once before simulating processes within the porous materials, including the various biological dynamics scenarios.

It is relatively straightforward to illustrate graphically on 2D cross sections how the resulting partitioning looks, when each partitioned pore region is depicted with a different color (Figure 2). However, the connections existing between the different regions are more evident when one zooms-in in 3D inside a portion of the image (Figure 3). This zooming-in highlights the compacity and the coherence of the pore regions.

**Figure 1.**
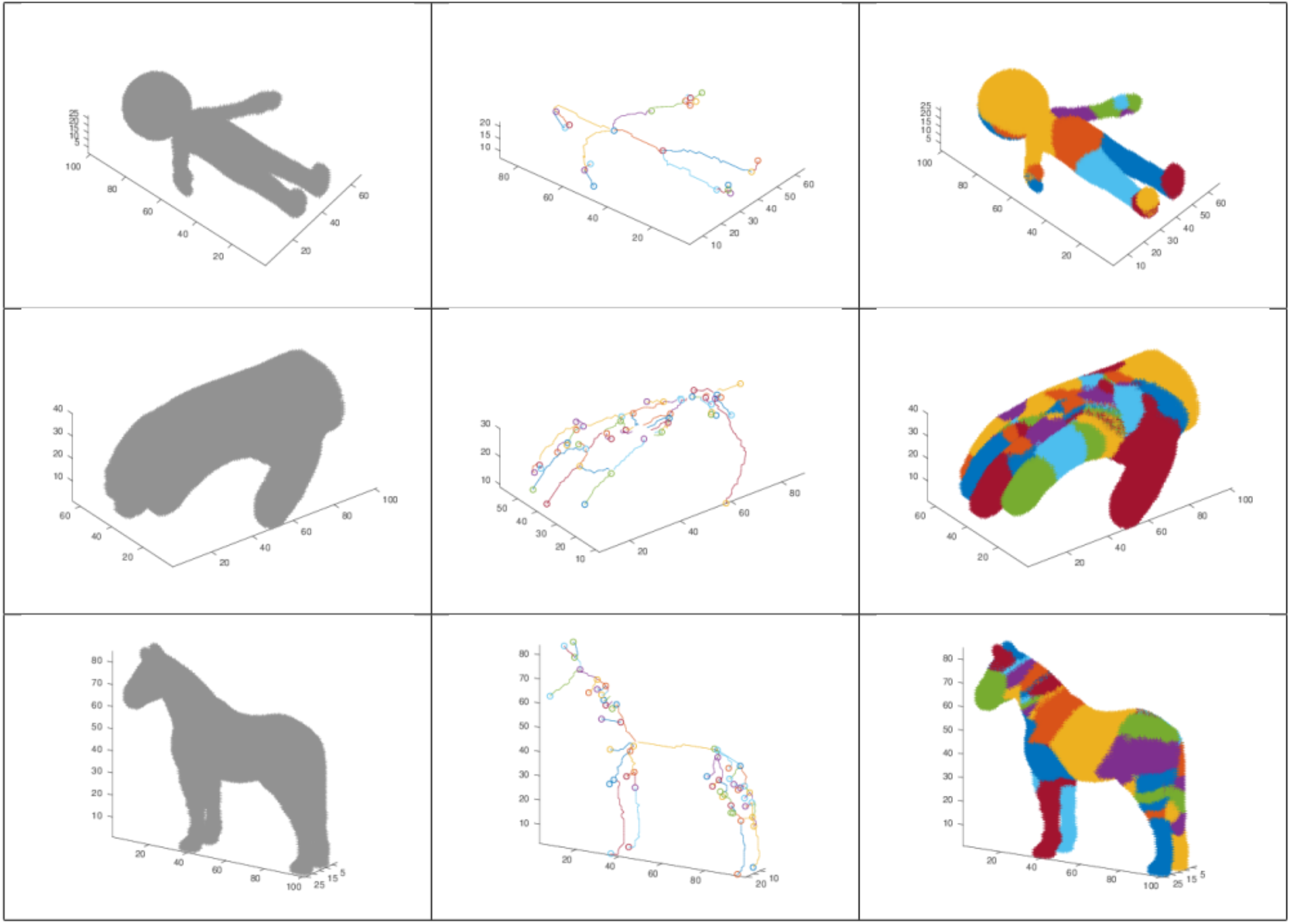
Illustrative examples of the partitioning of simple shapes (a doll, a hand, and a horse) using the curve skeleton. Left column: pictures of the voxelized shapes. Center: corresponding curve skeletons. Right: branch-based partitioning of the shapes. The calculations were carried out using the Matlab software. The 3D synthetic images are from the free 3D database Free3d (https://free3d.com/).

**Figure 2:**
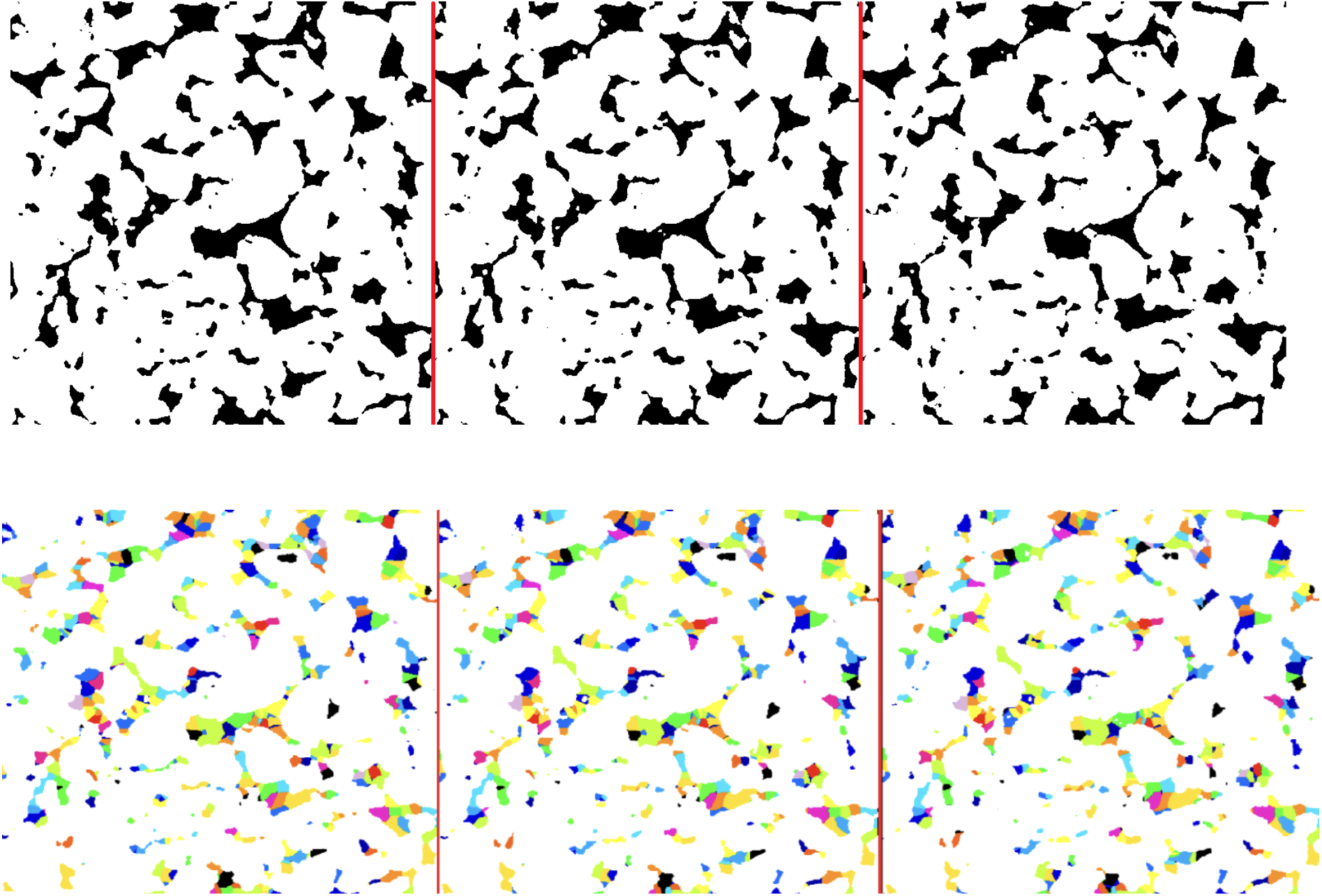
Top: three successive 2D cross-sections of the sandy loam soil (512×512×512 voxels; 24*μm*×24*μm*×24*μm*) where the voxels associated with the pore space are colored in black (17% of the whole image). Bottom: Corresponding partitioning of the pore space based on the curve skeleton

**Figure 3:**
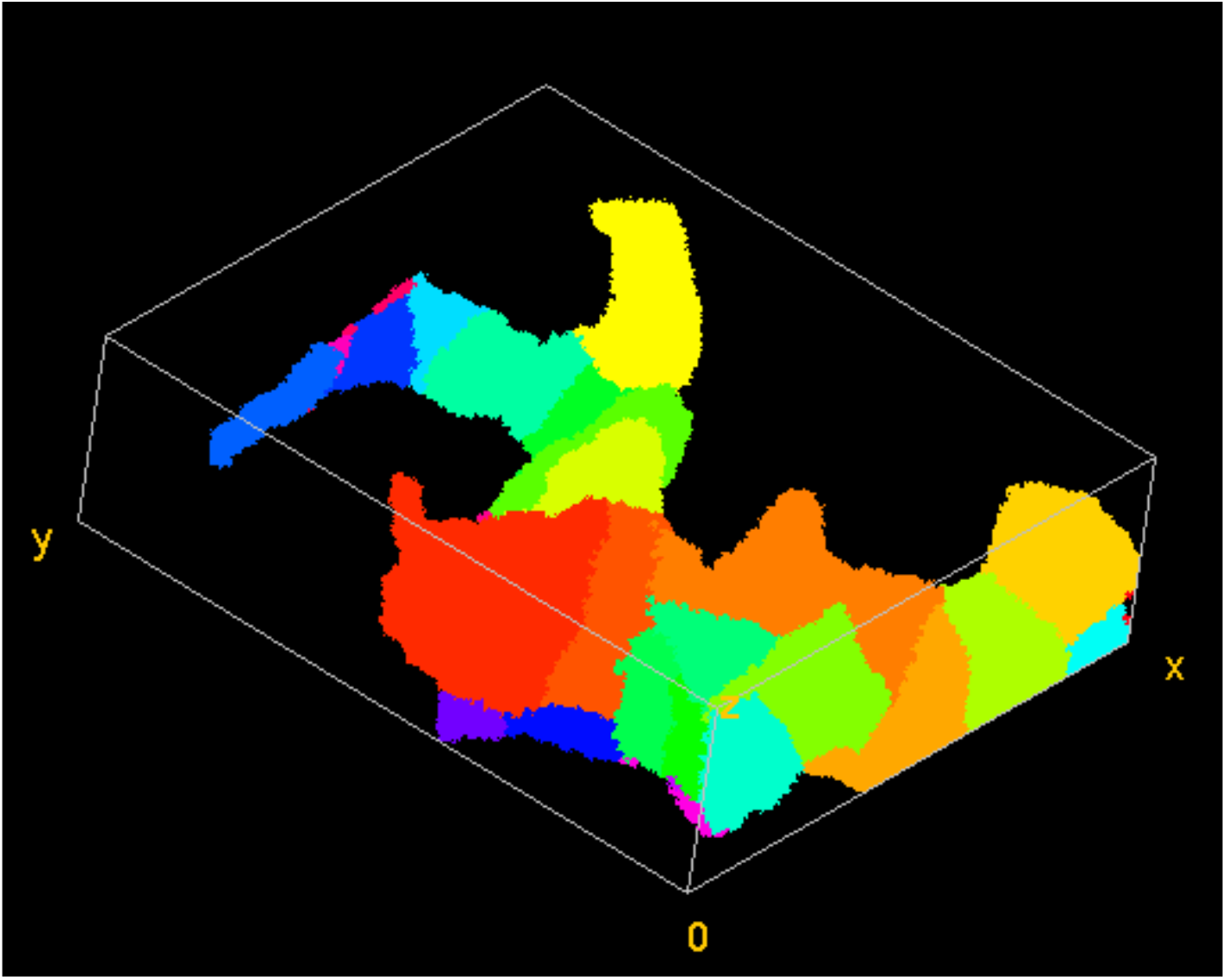
A zoom-in of the 3D projection of a portion of the partitioned image depicted in Figure 2. The 3D projection shows better than the 2D images (figure 2) how the partitioned pore regions are connected.

### 3.2) Simulation of diffusion processes

In order to use our geometrical representation for the (numerical) simulation of diffusion processes, we calibrate it by comparison with other approaches. We have implemented the same calibration process as the one described in Monga et al. [23]. Here, the goal of the calibration phase is to define the value of the diffusive overall conductance *∝*_*i,j*_.

To carry out this calibration, we introduce a given mass of organic matter into the first two planes. We compare the simulation results of the diffusion process through different cross-sections within the system using the three geometrical representations of the pore space: a voxel-based (LBM), ball-based (MOSAIC), and region-based (approach of the present paper). For the voxel-based approach, the Lattice Boltz-mann Method was used to simulate diffusion processes. For the balls- and regions-based approaches, as explained below, graph updating allows to simulate diffusion. Comparison of the different depth-concentration profiles obtained after adjustment of the conductance with the region-based approach with those resulting from the Mosaic-based approach (itself already compared in the past with the LBM method) suggests that an optimal fit of these different methods is obtained by setting *∝*_*i,j*_ to a constant value, equal to 0.35 (Figure 4)

**Figure 4:**
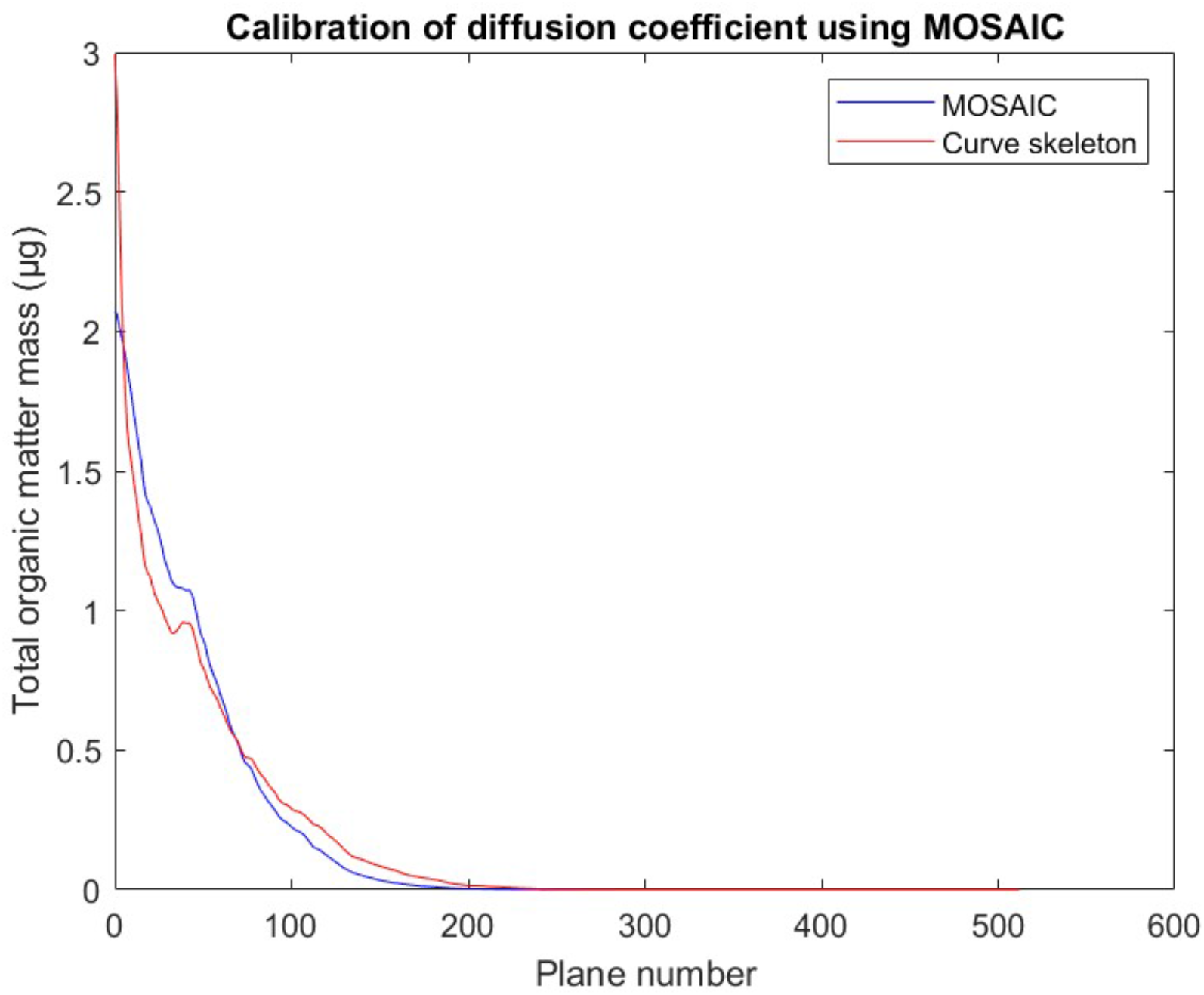
Example of the type of simulated results (for data set 1) used to calibrate the overall conductance to be used in later simulations. The X-axis displays the number of planes within the image (300 planes in total), whereas the Y-axis displays the total mass of organic matter within each plane. At the start, 592,7593 mg of carbon were introduced within the first two planes. The total simulation time was 1.783 hours. The optimal value of the diffusive over-all conductance coefficient *∝*_*i,j*_ was determined to be equal to 0.35, with an intercorrelation of 0.9818.

The computational cost of the diffusion phase is roughly proportional to the number of nodes of the geometrical primitive graph. For data set 1, using the geometrical modelling scheme described by Monga et al. [23], 191,583 balls are involved in the description of the pore space. The use of the curve skeleton-based algorithm described here results in the identification of 18,508 regions. Thus, the computing time for the diffusion simulation is divided by a ratio of approximately ten. For dataset 3, we obtained 478,191 balls and 72,691 regions, yielding a ratio of 7.

### 3.3) Simulation of microbial mineralization

The microbial mineralization of organic matter is described as the second step of a two steps sequence, the first step involving the diffusion of dissolved organic molecules, which is described using an implicit numerical scheme.

For dataset 1, we compare the results obtained by using the skeleton-based pore network graph with the ones obtained under identical conditions in an earlier publication [23] via the use of LBM and MOSAIC (Figure 5). Although there are some small numerical differences between the curves obtained with these different methods, their overall similarity is particularly noteworthy, given the large differences in computing times they require. On the personal computer that we used for the calculations, the computing time (CPU) for the LBM approach was several days; the Mosaic ball-based approach took 20 minutes, and the new method, based on partitioning the pore space with the curve skeleton, yielded its result in one minute.

**Figure 5:**
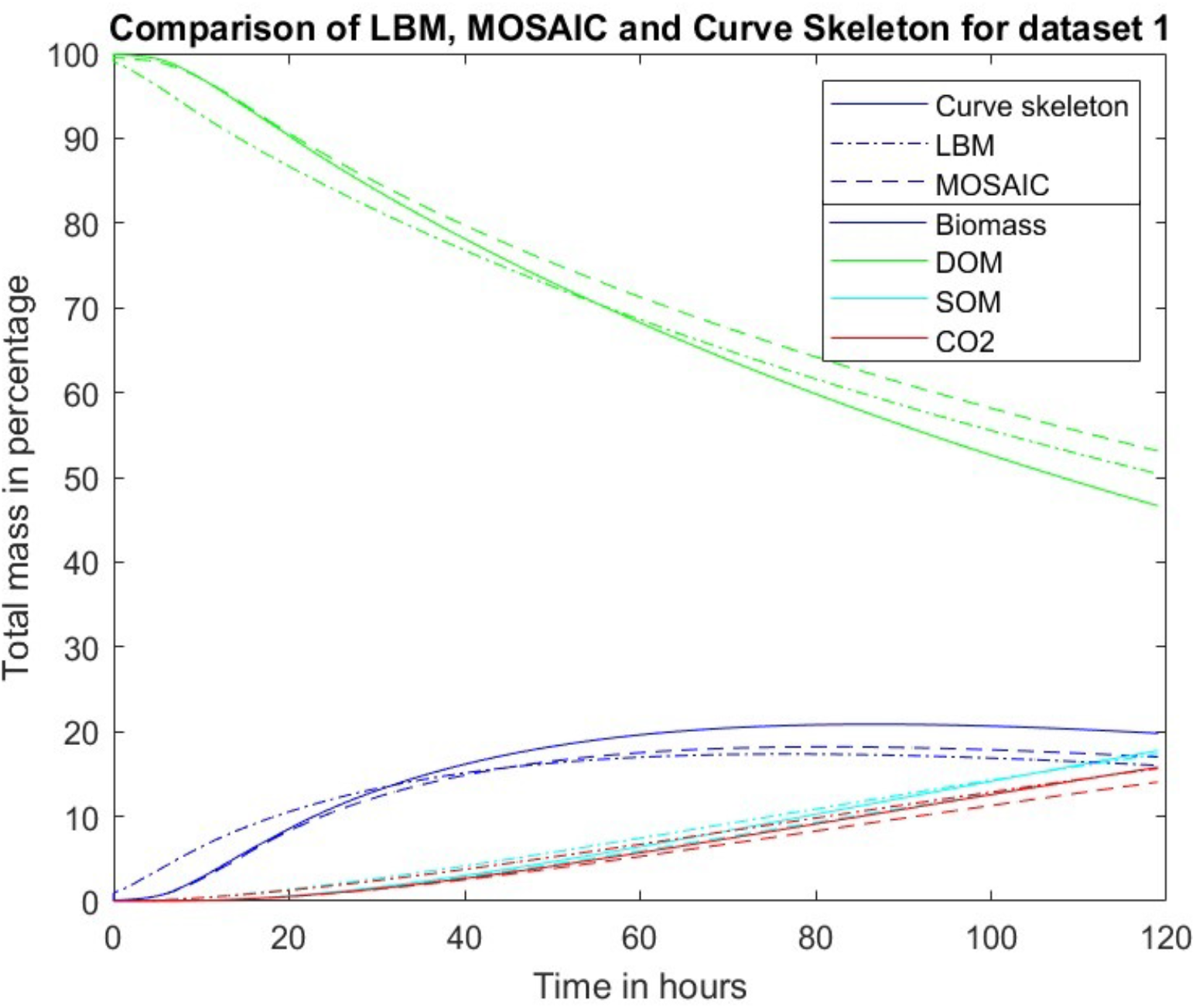
Comparison of different simulations of microbial mineralization for dataset 1 over a total simulation time of 5 days. The Y axis displays the mass of several components of the system expressed as a percentage of the total initial carbon mass. Simulations were carried out in the “curve skeleton” method with a diffusive conductance coefficient equal to 0.35.

For datasets 2,3, and 4, comparison was carried out simply with MOSAIC, involving a network of spherical balls, since the latter has already been compared favorably to simulations with the LBM method [53]. The biological parameters and the initial masses of micro-organisms and DOM were the same as for dataset 1, and the Dissolved Organic Matter (0. 2895 mgC) was likewise, at first, distributed homogeneously in the pore space. We put two spots of micro-organisms in the two biggest balls, with a radius of 27 and 30, respectively, considered in the Mosaic ball-based simulations. These balls correspond roughly to the volume of the 10 biggest regions when using the curve skeleton-based approach. Therefore, to enable a comparison between the two different simulation methods, we placed in these regions the same biomass as in the two largest balls in the Mosaic simulations.

Simulation results for data set 2 (Figure 6) show that the different curves are virtually indistinguishable for the two methods, in spite of considerable differences in the time required for computation. The simulations took 10 hours of CPU time with the Mosaic approach, versus merely one hour with the curve skeleton-based model. The results in Figure 6 also demonstrate that the diffusive overall conductance coefficient does not need (at least for these datasets) to be updated when changing datasets. Similar agreements between the two simulation methods were obtained for data sets 3 and 4 (data not shown), as well as a comparable 10-fold speeding up of the time needed for completion.

**Figure 6:**
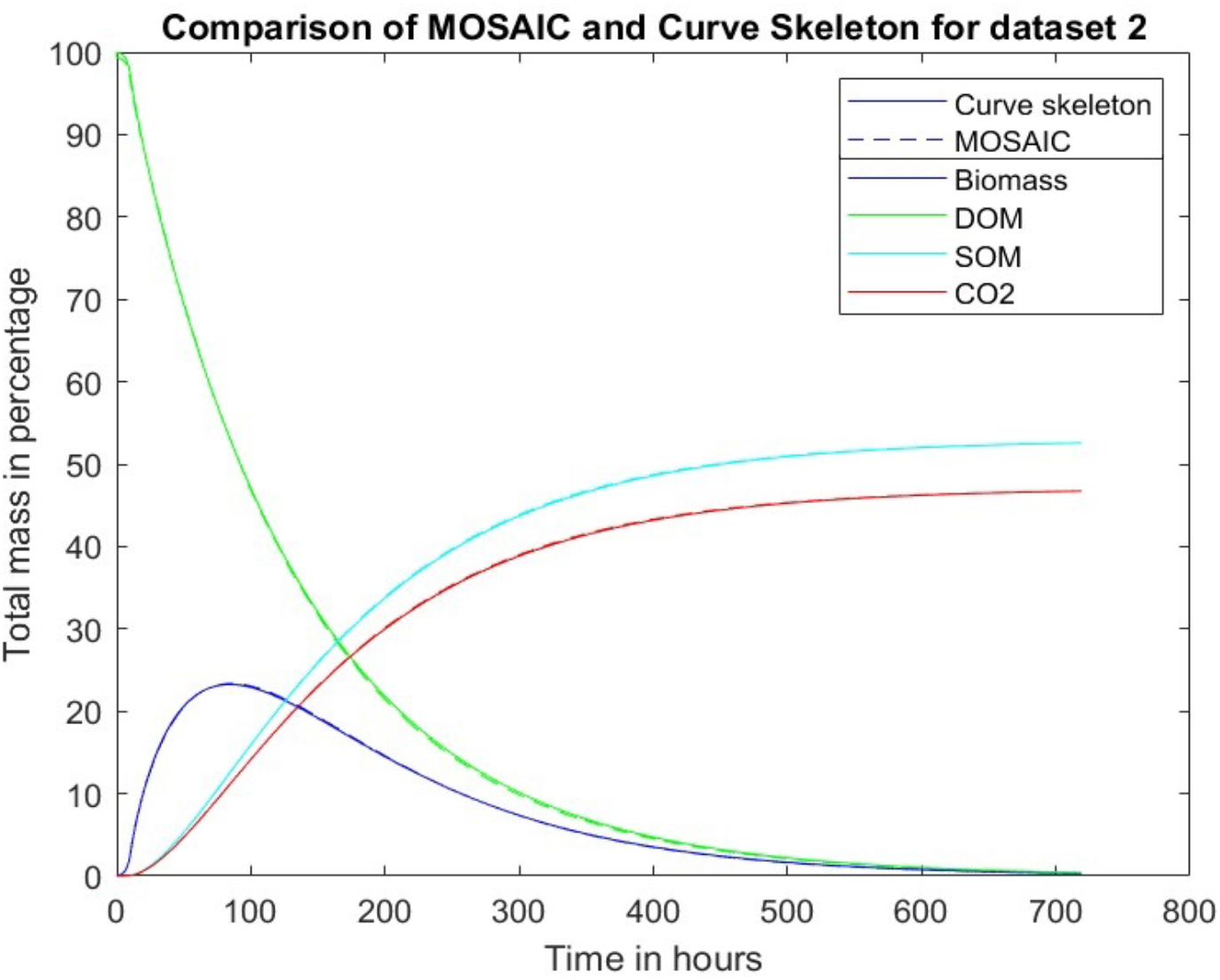
Comparison of simulations of microbial mineralization for dataset 2 using the ball-based MOSAIC program and the novel approach introduced here, based on the partitioning of the pore voxels on the basis of the curve skeleton. The simulations extended over 30 days and were carried out in the “curve skeleton” method with a diffusive conductance coefficient equal to 0.35.

## 4 Conclusion

The key result of the present article is the demonstration that the use of the curve skeleton to partition the pore space in 3D micro CT (Computed Tomography) images of soil samples, followed by the use of this partition for the simulation of the microbial mineralization of soil organic matter, leads to considerably shorter computing times. As far as we are aware, this use of the curve skeleton to represent complex volume shapes is the first in the context of soils. The method has been used in other disciplines in the past, like medicine, material sciences, chemical engineering, and in the analysis of porous media (e.g., carbonate rocks) but never to speed up the otherwise extremely lengthy simulations needed to describe the fate of organic matter in soils subjected to environmental change.

The principle of our method consists of segmenting the curve skeleton into simple branches, and afterward attaching to each simple branch a connected set of points. The result is a partition of the pore space, which can be represented by an attributed relational graph. In this graph, each node corresponds to a pore, and each arc to an adjacency between a couple of pores. We show that this geometrical representation of the pore space can be used directly to simulate microbial mineralization dynamics including diffusion and transformation processes. We assess the soundness of the approach by comparison with other methods that have been developed and used in the past in the same context. A possible drawback of the novel approach, compared with geometrical primitives-based modeling, is that each pore is defined by a set of connected voxels with no explicit geometric properties. On the other hand, the advantages of the pore space modeling based on the curve skeleton are fourfold. It does not impose specific shapes for a pore as do primitive-based modeling methods, involving balls or ellipsoids. It defines an exact piece-wise representation of the pore space, without losing any part of it, since it is based strictly on a partitioning of the pore space into distinct regions. When implementing diffusion processes, it involves an exact representation of the contact surfaces between adjacent regions. Finally, the number of nodes (pores) is much less than the ball network used in a previous work. To maximize the advantages and minimize the drawbacks, it might be possible in the future to implement hybrid geometric modeling algorithms using both skeletonization and geometric primitives.

## 5 Data

We use four different data sets in the simulations (Table 1). The first, already used by Monga et al. [23], consists of an image of a sandy loam soil (sand, silt, clay: 71, 19 and 10% soil mass, respectively) from the Bullion Field, an experimental site situated at the James Hutton Institute in Invergowrie (Scotland). A detailed description of the soil samples and of the methods used to produce CT images can be found in [54]. Data sets 2 and 3 consists of two CT images of quartz sand obtained in the Fontainebleau forest, corresponding respectively to high and low porosities. The size of these 3D CT images is 512×512×512 and the resolution 4.5*μm*×4.5*μm*×4.5*μm*, and their porosities are 57% and 42% respectively [25]. The fourth dataset is a Carbonado Diamond downloaded from the Digital Rocks portal. It is found only in placer deposits and Mesoproterozoic meta conglomerates in Bahia, Brazil [55].

**Table 1:** Comparative table about the characteristics of the different datasets.

|  | Dataset 1 | Dataset 2 | Dataset 3 | Dataset 4 |
| --- | --- | --- | --- | --- |
| Type | Sandy loam soil | High porosity quartz sand | Low porosity quartz sand | Porous polycrystalline diamond |
| Dimensions | 512 x 512 x 512 | 512 x 512 x 512 | 512 x 512 x 512 | 480 x 480 x 480 |
| Resolution | 24 $\mu\text{m}$ | 4.5 $\mu\text{m}$ | 4.5 $\mu\text{m}$ | 1 $\mu\text{m}$ |
| Pore space voxels | 22,720,090 | 74,766,570 | 57,350,410 | 27,669,358 |
| Pore space ratio | 17% | 57 % | 42% | 25 % |
| Number of regions | 18,508 | 259,299 | 68,090 | 45,386 |
| Number of balls | 191,583 | 23,762,616 | 478,191 | 407,055 |

## 6 Methods and approaches

From the 3D CT images, we extract a voxel-based representation of the pore space, typically encompassing hundreds of millions of voxels (Figure 2, top). In the first step, we compute the curve skeleton of this set of voxels [56]. In the second step, we segment the curve skeleton into maximal simple branches. In the third step, we identify for each voxel the simple branch that is the closest in an Euclidean distance sense. Afterward, we attach to each simple branch the corresponding set of voxels. In the fourth step, we label (i.e., partition) the pore space into connected sets of voxels (regions). In the fifth step, we build an attributed adjacency graph of regions from the partition where each node is attached to a set of connected voxels corresponding to a pore. Finally, we use this compact pore space representation to carry out the numerical simulation of microbial mineralization.

### 6.1. Geometrical modelling of the pore space using curve skeleton

#### 6.1.1 Computation of the curve skeleton

Let S be the set of voxels forming the pore space. Let B be the surface border of S. We assume without loss of generality that S is connected. If it is not the case, we work separately on each connected component of S. We compute the curve skeleton of S by means of successive erosions using the homotopic thinning algorithm.

As mentioned in [56,57], at each stage of the erosion process, voxel (i,j,k) is deleted if the following conditions hold :

- (i,j,k) lies on B (S boundary), i.e., at least one of its 26 neighbors lies outside S.
- (i,j,k) is not an ending point (an ending point has only one neighbor), regarding the 26-neighborhood.
- (i,j,k) removal does not modify Euler characteristics (topological condition).
- (i,j,k) removal does not modify the number of connected components (topological condition).

This erosion process (thinning) is iterated until the fixed point is reached (idempotence). The homotopic thinning algorithm is robust and works for any shape represented by voxels, even for disconnected B with connected S (volume shape with holes). By construction, the curve skeleton thickness is one voxel. In general, the curve skeleton of a given shape is unique, but some very symmetric shapes can have several curve skeletons, for instance the cube [0,9]^3^minus the concentric cube [3,6]^3^. For these cases, homotopic thinning algorithm comes up with a curve skeleton depending on implementation details, like the order in which boundary voxels of S(B) are considered. Such cases are unlikely when modelling natural shapes such as soils.

#### 6.1.2 Extraction of the pores: Partitioning of the curve skeleton into simple branches

We propose to segment the curve skeleton in order to extract the pore network.

The curve skeleton is represented by a set of voxels connected. Neighboring voxels are linked by an edge, we use the 26-connectivity.

In the curve skeleton, there are three different kinds of voxels:

- Ending voxels are nodes with only one neighbor.
- Simple voxels are nodes with exactly two neighbors.
- Interior voxels are nodes with more than two neighbors.

A branch is a maximal set of connected simple voxels. Ending voxels and interior voxels are located at the extremity of branches. Interior voxels lie at the interconnection between branches. Figure 1 shows an example of skeleton representation. We segment it into a set of branches using a straightforward algorithm. Thus, we get a set of branches forming a partition of the curve skeleton.

### 6.2) Simulation of microbial mineralization of organic matter in the extracted pores using graph-based representation

Afterwards, each voxel of S is associated with the nearest branch of the curve skeleton in the sense of Euclidean distance. The result is a partition of S composed of regions forming a practically connected set of voxels. In very uncommon cases, where the set of voxels attached to a branch would not form a connected component, one can split it into connected sets. Another way of tackling this problem would be to use the geodesic distance instead of the Euclidean one, as proposed in [58]. This way, the pore space can be partitioned into connected sets of voxels corresponding to a single branch of the curve skeleton. The partitioning of S is directly defined by the nearest sets attached to the branches. Three examples of volume shape partitioning of familiar shapes are given in Figure 1. At the end, we get an attributed adjacency graph representing the pore network denoted as G. The graph of regions defining the pore network is built as follows. Each node of the graph is attached to a region (pore) corresponding to a branch of the curve skeleton. Each arc of G is attached to a pair of neighboring regions.

The adjacency relationships are computed by building a 3D image L(i,j,k) where each voxel is given the label of the region where it is included. In case (i,j,k) does not belong to S, L(i,j,k) is set to 0. The arcs of G and the area of the contact surfaces between adjacent regions (pores) are determined by means of a one-pass scanning. For each voxel (i,j,k) of S we look for the three neighbors: (i+1,j,k), (i,j+1,k), (i,j,k+1) and for each neighbor belonging to S, we update the graph adjacencies (arcs) and the areas of the contact surfaces (arc features) by looking to the values of the label image (L). If the two labels are the same, we do nothing. If the two labels are different, we eventually create a new arc in the graph and increment the area of the contact surfaces between the two regions (Figure 7).

**Figure 7:**
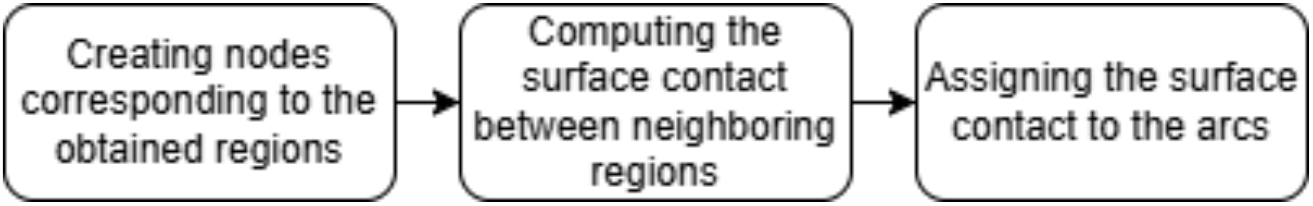
Schematic representation of the main steps involved in creating the graph of segmented regions.

This algorithm finds all neighboring relationships between regions and all corresponding areas of contact surfaces. It is used because not all edges of G can be created considering only the branches of the curve skeleton. Indeed, some regions may share a common boundary surface although their branches do not have a common interior node. Neglecting regions adjacencies, as was done in [41], could be a significant drawback for many applications. For instance, in the case of diffusion simulations, it would imply that flows would transit only along branches of the curve skeleton.

#### 6.2.1 Simulation framework: Microbial mineralization of organic matter in the graph of regions

From the previous stages, we get an attributed relational graph G such that:

- Each node is attached to a connected set of voxels (region, pore) corresponding to a branch of the curve skeleton; we associate to each node the coordinates of the inertia center of the set of voxels and its volume (number of voxels).
- Each arc is attached to an adjacency relationship between two regions (pores); we associate to each arc the area of the contact surface between the two corresponding regions (pores).

Afterward, we use the graph G, representing the pore space, to simulate microbial mineralization. The global scheme of the numerical simulation is the same as the one described in [23]. However, the geometrical representation of the pore space is different. These authors described the pore space by the minimal set of balls recovering the surface skeleton. The drawback of that approach is that a part of the pore space (typically 15%) is lost due to piecewise approximation errors. The geometrical representation adopted in the present article covers completely the set of voxels corresponding to the pore space. In particular, we get exact values for the contact surface areas between two pores.

We get the following implicit numerical scheme:

We note: 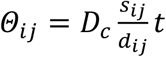 where *D*_*c*_, *s*_*ij*_, *d*_*ij*_, *t* are respectively the diffusion coefficient, the area of the contact surface between node (region, pore) i and j, the distance between the two inertia centers of regions i and j, and the discretization step time.

If we denote by *c*_*i*_, *v*_*i*_,*N*_*i*_, *ϑ* (*N*_*i*_), respectively, the concentration at node *i*, the volume of the node *i*, the node *i* and the neighbor of the node *N*_*i*_, the relationship between concentrations at successive iterations can be expressed as follows:

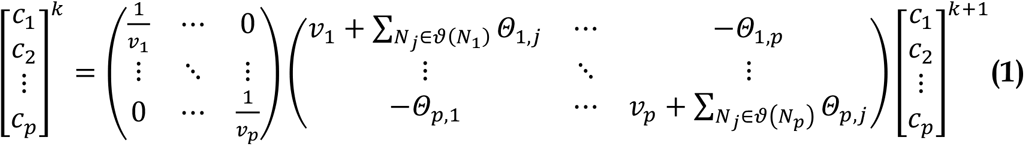

We solve the above system with help of the conjugate gradient method.

#### 6.2.2 Diffusive overall conductance computation

When the Fick flows are defined between two regions, we must multiply the flows with a coefficient *∝*_*i,j*_ called diffusive overall conductance such as in [59]. In order to define *∝*_*i,j*_, we calibrate by comparison with the diffusion simulation using the balls network and LBM. Practically, we find out that we can set *∝*_*i,j*_ to a constant value α.

Therefore the Fick flow becomes:

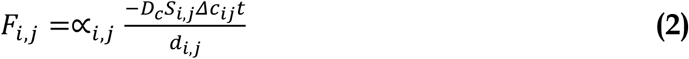

The calibration principle is to adjust such that the diffusion using the curve skeleton-based pore network description fits with the ones provided by the Lattice-Boltz-mann Model (LBM) and balls network methods. Practically, we define diffusion benchmarks and optimize the correlation between the outputs. In this way we find out that setting *∝*_*i,j*_ to the constant value 0.35 allows good fitting results. From experience, it turns out that this applies for different sets of data.

#### 6.2.3 Step time validation using an explicit numerical scheme

We can check the pertinence of the step time value for the implicit numerical scheme using the explicit numerical scheme. In a previous paper we proposed to implement an implicit numerical scheme using directly the graph of Monga et al. [23]. In the present work we use a matrix representation yielding substantial computational gain. With the same calculation scheme as in section 3.5.2, we get:

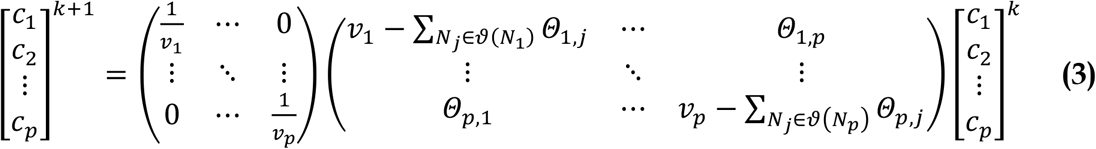

The drawback of the explicit numerical scheme is that it requires very small time steps to avoid negative values. Thus, we use it only to check the validity of the implicit numerical scheme’s time step.

#### 6.2.4 Modelling of microbial mineralization

The principle is to discretize the process in time and to successively implement transformation and diffusion processes. The diffusion processes are implemented by means of the implicit numerical scheme (see 3.5.2 and 3.5.3). The computational cost is roughly proportional to the number of graph nodes, which is the number of balls. In most cases, the number of regions provided by the curve skeleton-based method is much lower (ratio of 1/10 to 1/20).

To simulate the microbial mineralization of organic matter, the same biological parameters that Monga et al. [25] considered for *Arthrobacter sp. 9R* were taken for DOM degradation with 9.6 j^-1^ for the maximum growth rate, 0.001 gC g^-1^ for the constant of half-saturation, 0.2 j^-1^ for the respiration rate, 0.5 j^-1^ for the mortality rate and 0.55 for the portion of microbial biomass that returns to DOM. A rate of 0.3 j^-1^ was adopted for the mineralization of POM and one of 0.001 j^-1^ for the mineralization rate of SOM. We put 5.2 10^7^ bacteria (0.18 µg biomass) divided into 1000 spots in the pore space, and a mass of DOM of 0. 2895 mgC in the pore space corresponding to the concentration of 0.13 mgC/g soil. We use the molecular diffusion coefficient of DOM in water of 6.73 10-6 cm2 s^-1^.

## 7 Acknowledgments

The research described in the present article was made possible through CNRST grant I-Maroc (APRD program) and through project Microlarge financed by the Agence Nationale de la Recherche (Paris, France).

## Notes

### Competing Interest Statement

The authors have declared no competing interest.

